# Omission responses in field potentials but not spikes in rat auditory cortex

**DOI:** 10.1101/2022.02.11.479668

**Authors:** Ryszard Auksztulewicz, Vani Gurusamy Rajendran, Fei Peng, Jan Wilbert Hendrik Schnupp, Nicol Spencer Harper

## Abstract

Non-invasive recordings of gross neural activity in humans often show responses to omitted stimuli in steady trains of identical stimuli. This has been taken as evidence for the neural coding of prediction or prediction error. However, evidence for such omission responses from invasive recordings of cellular-scale responses in animal models is scarce. Here, we sought to characterise omission responses using extracellular recordings in the auditory cortex of anaesthetised rats. We profiled omission responses across local field potentials (LFP), analogue multiunit activity (AMUA), and single/multi-unit spiking activity, using stimuli that were fixed-rate trains of acoustic noise bursts where 5% of bursts were randomly omitted. Significant omission responses were observed in LFP and AMUA signals, but not in spiking activity. These omission responses had a lower amplitude and longer latency than burst-evoked sensory responses, and omission response amplitude increased as a function of the number of preceding bursts. Contrary to theories of neural entrainment, rhythmic stimulus presentation did not increase low-frequency phase-locking of neural activity specific to the stimulus presentation rate. Together, our findings show that omission responses are observed in LFP and AMUA signals, with laminar specificity, but are not observed in spiking activity, and do not show evidence for low-frequency phase locking. This has implications for models of cortical processing that require many neurons to encode prediction error in their spike output, and may have some consistency with representation of error in dendrites electrotonically distant from the soma.

## 1. INTRODUCTION

At least since the times of von Helmholtz (von Helmholtz, 1867), prediction has been proposed as important to perception, and many principled models of cortical function have prediction of current or future sensory input as a central component (Chalk et al., 2018; Friston, 2003; Olshausen and Field, 1996; Rao and Ballard, 1999; Singer et al., 2018). Efficient prediction of future sensory inputs may facilitate action guidance and sensory feature extraction (Bialek et al., 2001), and hence may be a key principle governing sensory neural systems, arguably explaining many of their features (Singer et al., 2018). Thus finding neural representations of predictions, or of prediction error, the deviation of sensory inputs from their predictions, has been a recent area of intense research focus. Central to these investigations have been oddball paradigms, in which a sequence of expected stimuli is replaced by an unexpected stimulus (Carbajal and Malmierca, 2018; Garrido et al., 2009b); these paradigms have provided insights into neural prediction using behavioural (Simpson et al., 2014), EEG/MEG (Auksztulewicz and Friston, 2015; Garrido et al., 2009b; Näätänen et al., 2007), and *in vivo* neurophysiological measurements (An et al., 2020; Parras et al., 2017; Ulanovsky et al., 2003). However, rather than altering the expected stimulus, it can instead be omitted, enabling observations of the form and timing of predictive signals that are not confounded by processing of incoming stimuli (Schröger et al., 2015). Although omission responses have often been reported using measurements of neural activity in humans that are non-invasive (Bendixen et al., 2009; Chennu et al., 2016; Dercksen et al., 2020; Raij et al., 1997; Sanmiguel et al., 2013; Todorovic et al., 2011; Wacongne et al., 2012; Yabe et al., 1997) or from the cortical surface (Fonken et al., 2019; Hughes et al., 2001), there has been little investigation at a detailed level by using penetrating electrodes.

Non-invasive human studies suggest that responses to omitted stimuli peak later than stimulus-evoked responses and are often reported to have lower amplitudes (Andreou et al., 2015), but see (Horváth et al., 2010). Crucially, omission responses have a broad frequency spectrum (Todorovic et al., 2011) and are time-locked to the expected onset of an omitted stimulus (Chennu et al., 2016). This differentiates them from offset responses, which can also be evoked by interrupting a stimulus train but are time-locked to the last presented stimulus (Fig. 1A) (Chien et al., 2019), and from entrainment which is the hypothesised phase-alignment of ongoing low-frequency neural activity to the rhythmic structure of isochronous stimulus sequences (Haegens and Zion Golumbic, 2018; Lakatos et al., 2019). Earlier human studies have suggested that omission responses can be observed only for relatively fast stimulus presentation rates, well above 5 Hz (Yabe et al., 1997), which was interpreted as a limited window of temporal integration. However, mounting evidence demonstrates that omission responses can also be observed in awake humans following longer inter-stimulus intervals, in the sub-second (Fonken et al., 2019; Halgren et al., 1995) and supra-second range (Busse and Woldorff, 2003). Omission responses at different time scales may be differentially influenced by cognitive factors: for instance, omissions following short ISIs (periodicity > 5 Hz) have been suggested to be elicited entirely automatically, while slower time scales (periodicity < 2 Hz) may be modulated by attention (Chennu et al., 2016; Chien et al., 2019; Karamürsel and Bullock, 2000), but see (Hughes et al., 2001).

**Figure 1.**
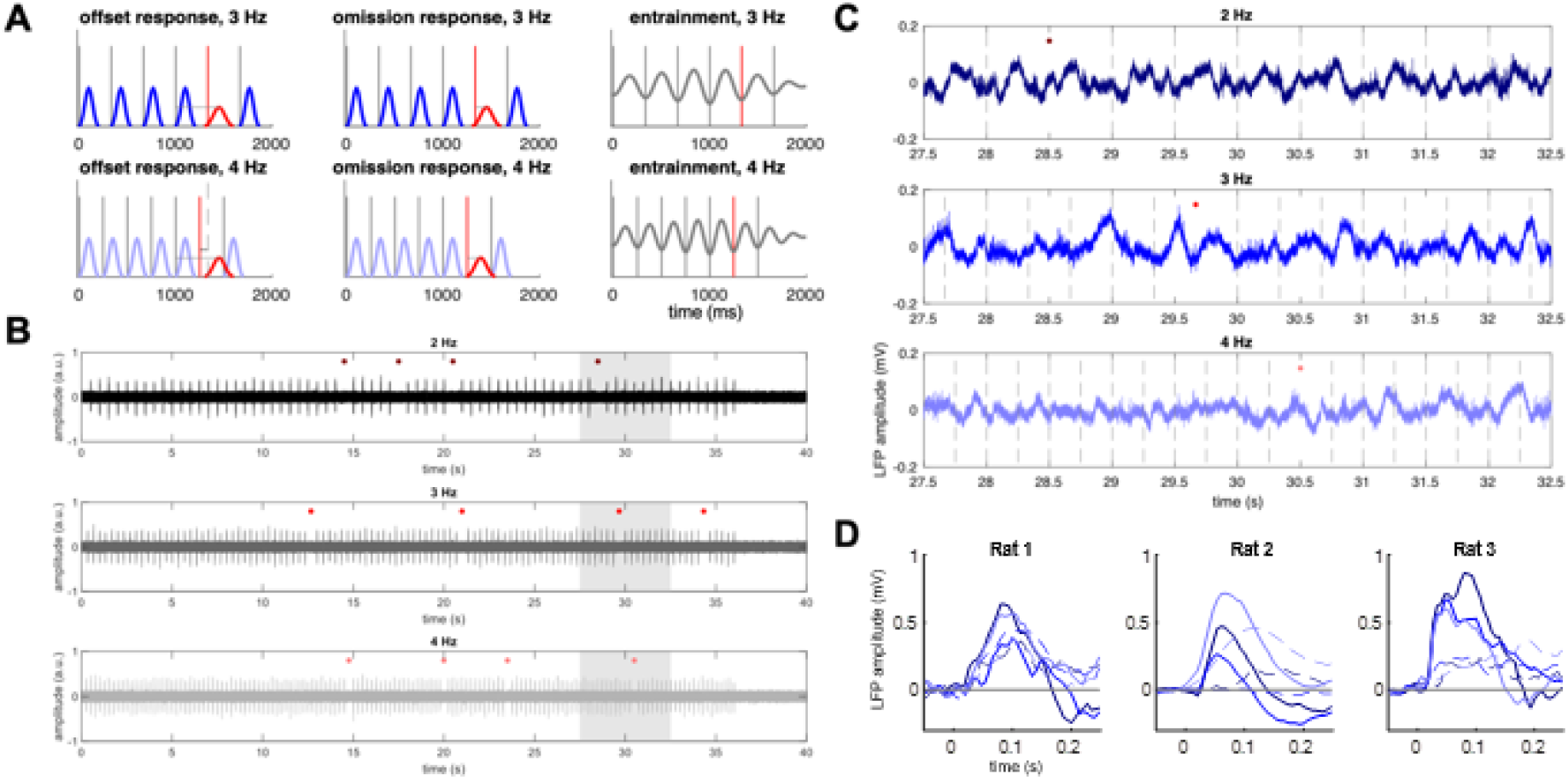
**(A)** Schematic representing three possible types of neural responses that could occur to omissions of rhythmically presented stimuli. Vertical lines represent stimuli presented at 3 Hz (dark grey lines, upper plots), 4 Hz (light grey lines, lower plots), or omitted (red lines). Hypothetical responses are plotted in blue. Left: offset response, whose latency is locked (grey horizontal line) to the preceding stimulus rather than to the omitted stimulus. Middle: true omission response, whose latency is locked (grey horizontal line) to the omitted stimulus rather than to the preceding stimulus. When analysing response latencies relative to the omitted stimulus, true omission responses should show the same latency for both 3 and 4 Hz, while offset responses should show a latency shift between 3 and 4 Hz (lower left plot, difference between solid red and dashed grey vertical lines). Right: low-frequency entrainment of ongoing neural activity to the stimulus sequence. **(B)** Example stimulus waveforms for 2 Hz (upper plot), 3 Hz (middle plot), and 4 Hz (lower plot) sequences. Filled circles denote omitted stimuli. Shaded area denotes the time segment for which raw local field potentials are plotted in (C). **(C)** Example local field potentials from a representative electrode. Dashed vertical lines denote presented stimuli. Filled circles denote omitted stimuli. **(D)** Examples of LFP responses, plotted for a representative channel for each rat. Solid lines: stimulus-evoked responses; dashed lines: omission responses; colours as above.

These findings in humans are seemingly inconsistent with invasive cellular-scale studies in animal models, where omission-related activity has been found in extracellular signals recorded in the auditory cortex of trained awake macaques attending to auditory streams presented at rates of ~2 Hz (Lakatos et al., 2013), but also in anaesthetised rodents exposed to very slow (< 0.5 Hz) isochronous sequences. In the latter case, omission-related activity has been reported in the non-lemniscal thalamus of guinea pigs using intracellular recordings (Gao et al., 2009) and in the auditory cortex of mice (Li et al., 2017; Wang et al., 2018), although necessitating a large number of preceding standards. However, the last two studies used calcium imaging, which is characterised by slowly decaying calcium signals, rather than direct electrophysiological recordings of neural activity. To disambiguate the neural mechanisms of omission responses at faster time scales typical for humans, an animal model has yet to be established, and it is largely unknown whether such an animal model can successfully replicate findings from humans, even in anaesthetised animals which are naive to the standards used.

Here, we used penetrating microelectrodes to record local neural population activity from the auditory cortex of anaesthetised rats to trains of noise bursts at 2, 3 or 4 Hz, with noise bursts occasionally omitted. We observed that the local field potentials and analogue-multiunit activity at the majority of recording sites showed a response just after the time point when a stimulus was expected but omitted. These omission responses had a fixed latency relative to expected onset regardless of stimulus rate, indicating true omission responses rather than offset responses. The omission signals increased as a function of the number of preceding noise bursts, suggesting that they might be modulated by the strength of previously formed predictions. However, such omission responses were not apparent in the single-unit or multi-unit responses. We analysed the spectral and laminar profile of the omission responses, to characterise omission responses with a precision that cannot be achieved by non-invasive human recordings. This analysis revealed that, in superficial and intermediate layers, omission responses were relatively stronger in higher frequency bands than in lower frequency bands. Finally, we also tested whether rhythmic stimulus presentation increased low-frequency phase-locking of neural activity specific to the stimulus presentation rate. However, contrary to theories of neural entrainment, no such low-frequency phase-locking was found. Together this suggests that prediction or prediction error might be represented in dendritic activity or input to auditory cortex, but its representation is rare or non-existent in spiking activity of auditory cortical neurons, at least under the conditions examined. This limits the space of possible models of cortex involving prediction.

## 2. RESULTS

### 2.1 Omission responses are present in LFP and AMUA signals, but not in single/multi-unit spiking activity

We recorded a total of 6 penetrations from 3 rats using multi-electrode eight-shank probes with 8 electrodes (channels) along each shank. Anaesthetised naive rats were exposed to trains of noise bursts presented at an isochronous rate of 2, 3 or 4 Hz, with a random subset of 5% of noise bursts omitted from each train (Fig. 1; see Materials and Methods). The probes penetrated perpendicularly through the auditory cortex to record across its full depth. This resulted in a total of 384 channels recorded. Data were analysed in two broad frequency bands, including lower (0.1-75 Hz; LFP analysis) and higher (300-6000 Hz; AMUA analysis) frequencies, which have been proposed to be predominantly sensitive to summed inputs and local outputs of neural populations respectively (Goense and Logothetis, 2008; Logothetis, 2002), but see (Burns et al., 2010). The data were additionally spike-sorted to yield 43 single units and 70 multi-units that passed a response reliability criterion (see Materials and Methods). Population LFP, AMUA, and spiking activity following presented and omitted bursts are shown in Fig. 2.

**Figure 2.**
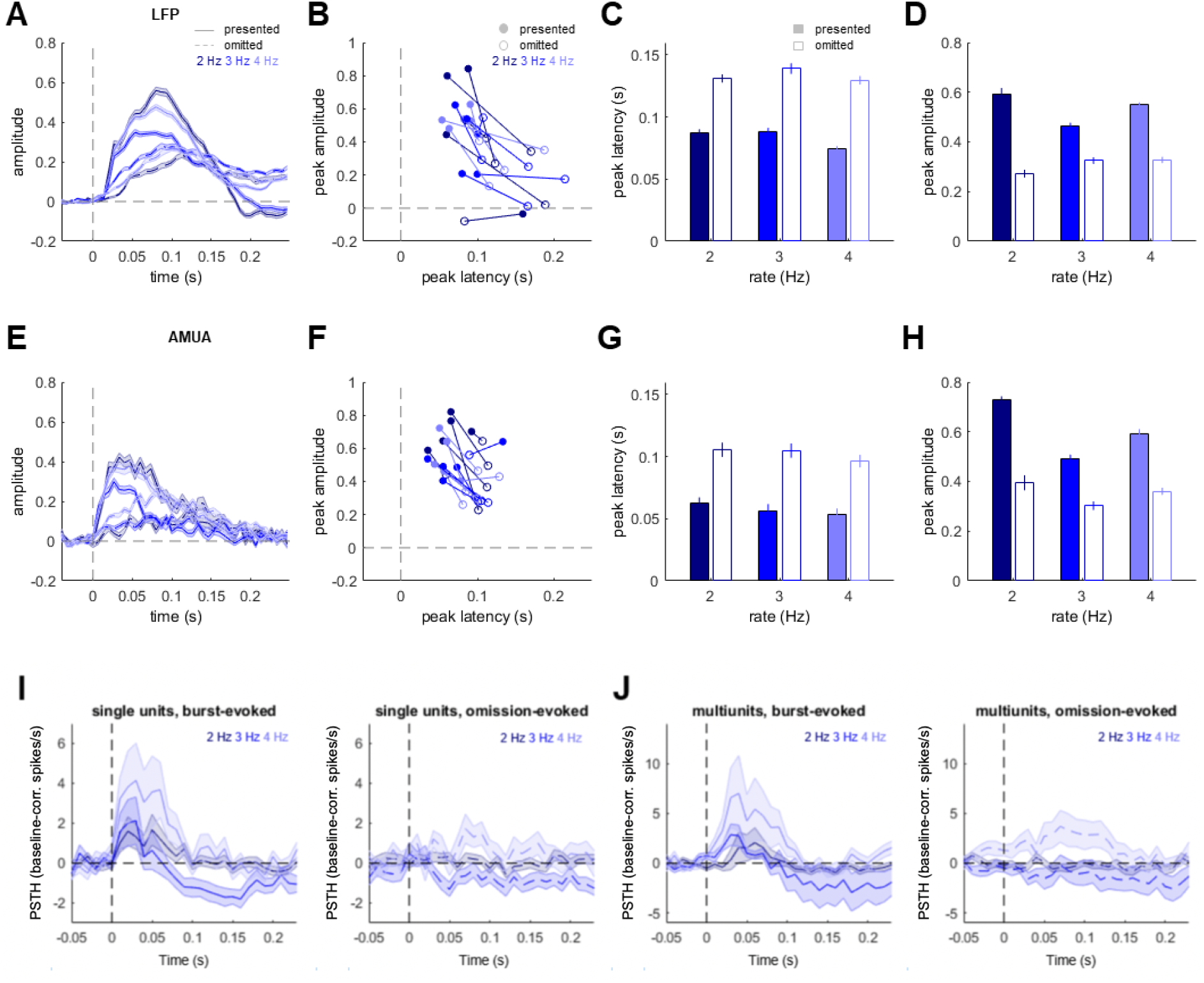
**(A - D)** Local field potential (LFP) responses to presented vs. omitted stimuli. **(E - H)** Analog multiunit activity (AMUA) responses to presented vs. omitted stimuli. **(A, E)** Time courses of responses (normalised to pre-stimulus baseline) evoked by presented stimuli (solid lines) relative to stimulus presentation, and by stimulus omissions (dashed lines) relative to the expected but omitted stimulus onset. Dark blue, blue, and light bluelines correspond to stimulus presentation rates of 2, 3, and 4 Hz. Filled black circles represent significant main effects of stimulus (presented vs. omitted; p < 0.05, FDR-corrected across time points). Shaded areas represent standard error of the mean (SEM) across channels. **(B, F)** Peak amplitudes (Y axis) and latencies (X axis) of each penetration, averaged across analysed channels. Filled circles: stimulus-evoked responses; empty circles: omission responses; colours as above. **(C, G)** Peak amplitude comparison of stimulus-evoked (filled bars) and omission responses (empty bars) across the three stimulus presentation rates. Error bars represent SEM across channels. Please note that single channels are presented in (B,F). **(D, H)** Peak amplitude comparison of stimulus-evoked vs. omission responses across the three stimulus presentation rates (filled/empty bars as above). Error bars represent SEM across channels. **(I)** Baseline-corrected peristimulus time histograms (PSTHs), quantifying single-unit spiking activity responses to noise bursts (left panel) and omitted stimuli (right panel). Colours as in (A, E). Shaded areas represent SEM across units. **(J)** Multiunit PSTHs. Legend as in (I).

First, to quantify the proportion of channels showing significant activity in the stimulus-evoked time window (0-250 ms relative to burst onset) or in the omission time window (0-250 ms relative to expected but omitted burst onset), for each channel we concatenated single-trial time-average amplitude estimates across the three burst rates and entered them into one-sample t-tests. In the LFP analysis (Fig. 2A), on average, 35.16 channels (SEM 9.60, corresponding to 54.95% ± 15% channels) per penetration showed significant responses to both presented sounds and sound omissions (averaged across rates), while in the AMUA analysis (Fig. 2E), on average, 21.33 channels (SEM 4.82, corresponding to 33.33% ± 7.53% channels) per penetration showed significant responses to both presented sounds and sound omissions (one-sample t-tests; in both LFP and AMUA, p_FDR_ < 0.05, false discovery rate corrected; (Benjamini and Hochberg, 1995)). Additionally, a number of channels responded only to presented sounds (LFP: 24.74% ± 15.26%; AMUA: 38.54% ± 9.06%), and a smaller proportion of channels responded only to omitted sounds (LFP: 9.64% ± 7.86%; AMUA: 4.16% ± 1.83%). No responses to presented or omitted sounds were recorded in the remaining channels (LFP: 7.81% ± 3.56%; AMUA: 25.26% ± 8.91%).

In contrast to LFP and AMUA signals, the analysis of spiking activity in 113 single and multi unit responses did not yield consistent omission responses. Among our 43 single units (with 11 units assigned to superficial layers, 10 to intermediate layers, and 22 to deep layers; see Materials and Methods), none showed a significant omission response while correcting for multiple comparisons (all p_FDR_ > 0.05; Fig. 2I), with only 3/43 units showing post-omission PSTH higher than the null distribution at an uncorrected p < 0.05. Similarly, among 70 analysed multiunits (with 18 units assigned to superficial layers, 15 to intermediate layers, and 37 to deep layers), none showed a significant omission response (all p_FDR_ > 0.05; Fig. 2J), and 9 units had the post-omission PSTH survive the uncorrected threshold of p < 0.05. While a visual inspection of both single- and multiunit activity did indicate relatively robust activity in the post-omission period for the 4 Hz burst rate, this activity started before the expected (but omitted) burst, suggesting that it corresponds to an train-offset response rather than to a true omission response. No such activity was observed for the slower burst rates (2 and 3 Hz).

### 2.2 Amplitudes and latencies of omission-evoked responses

Since we only observed omission responses in the LFP and AMUA data, we further analysed these two signal types. Our first aim was to compare the latencies and amplitudes of neural responses to presented and omitted sounds (Fig 2B, F), and the analyses that follow are thus focused on channels displaying significant burst-evoked and omission-evoked responses. In both LFP and AMUA analyses, single-channel data (peak amplitude or peak latency values) were entered into a mixed-effects model with two fixed-effects factors (stimulus type: burst vs. omission; presentation rate: 2, 3, and 4 Hz) and one random-effects factor (penetration). In both LFP and AMUA signals, there was a significant difference between responses evoked by burst omissions and preceding burst presentations in terms of the peak amplitude (main effect of stimulus type; LFP: F_1,1182_ = 10.14, p = 0.033, Fig. 2C; AMUA: F_1,732_ = 41.86, p < 0.001, Fig. 2G), with omission-evoked responses having lower peak amplitudes than burst-evoked responses. Additionally, in the AMUA but not the LFP, there was a trend towards a main effect of presentation rate (F_2,732_ = 3.8, p = 0.0563). However, the interaction between presentation rate (2, 3, and 4 Hz) and stimulus (omission vs. burst) was not significant in either LFP or AMUA analysis (both p > 0.18), suggesting that the relative strength of omission responses does not depend on presentation rate.

There was also a significant difference in peak latency between responses evoked by omitted and presented stimuli in LFP and AMUA (LFP: F_1,1182_ = 13.31, p = 0.021, Fig. 2D; AMUA: F_1,732_ = 38.21, p < 0.001, Fig. 2H), with omission responses peaking later than burst-evoked responses. Crucially, neither the main effect of presentation rate or the interaction between presentation rate and stimulus were significant (both AMUA and LFP analysis: p > 0.3), suggesting that the latency of omission responses is locked to the onset of an expected but omitted noise burst, and is not modulated by presentation rate.

The same pattern of results was replicated in a control analysis, in which all channels were included (rather than only those showing a significant response to bursts and omissions; Fig. S1). In the LFP data, both amplitude and latency differed between burst-evoked and omission-evoked responses (amplitude: F_1,1896_ = 8.99, p = 0.03; latency: F_1,1896_ = 10.9, p = 0.021) but the main effects of presentation rate and the interaction effects between presentation rate and stimulus were not significant (p > 0.15). In the AMUA data, beyond the main effect of stimulus on both amplitude (F(1,2232)=36.612, p = 0.002) and latency (F_1,2232_ = 60.35, p < 0.001), we also found a significant main effect of presentation rate on amplitude (F_2,2232_ = 26.29, p < 0.001) but not on latency (p > 0.3). The interaction effects between presentation rate and stimulus were not significant (p > 0.3).

A further control analysis was performed (Fig. S2) to confirm that the omission responses identified above were not due to random noise fluctuations in the post-omission time window. To this end, we shuffled single-trial responses over time points and analysed the data in the same manner as in the main analysis. The analysis of time-shuffled LFP data revealed no main effect of stimulus on signal amplitude (p = 0.0838) and no interaction between stimulus and presentation rate (p = 0.6405), but a main effect of presentation rate (F_2,2232_ = 580.57, p<0.001). The analysis of LFP response latency revealed no main or interaction effects (p > 0.5). Similarly, the analysis of time-shuffled AMUAdata revealed no main effect of stimulus on signal amplitude (p = 0.2104) and no interaction between stimulus and presentation rate (p = 0.2974), but a main effect of presentation rate (F_2,2232_ = 35.25, p<0.001). The analysis of AMUA response latency revealed no main or interaction effects (p > 0.25).

To test whether omissions can be attributed to the same channels which show evoked responses, we analysed the correlations between the peak amplitudes of evoked and omitted responses across channels. This analysis revealed significant correlations for both types of analyses (LFP, AMUA) and all presentation rates (2, 3, and 4 Hz). In the LFP analysis (Fig. 3A), the correlation coefficients were r = 0.4465, p < 0.001 (2 Hz); r = 0.2185, p = 0.002 (3 Hz); r = 0.5401, p < 0.001 (4 Hz). In the AMUA analysis (Fig. 3B), the correlation coefficients were r = 0.1515, p = 0.0433 (2 Hz); r = 0.1947, p = 0.0091 (3 Hz); r = 0.4367, p < 0.001 (4 Hz).

**Figure 3.**
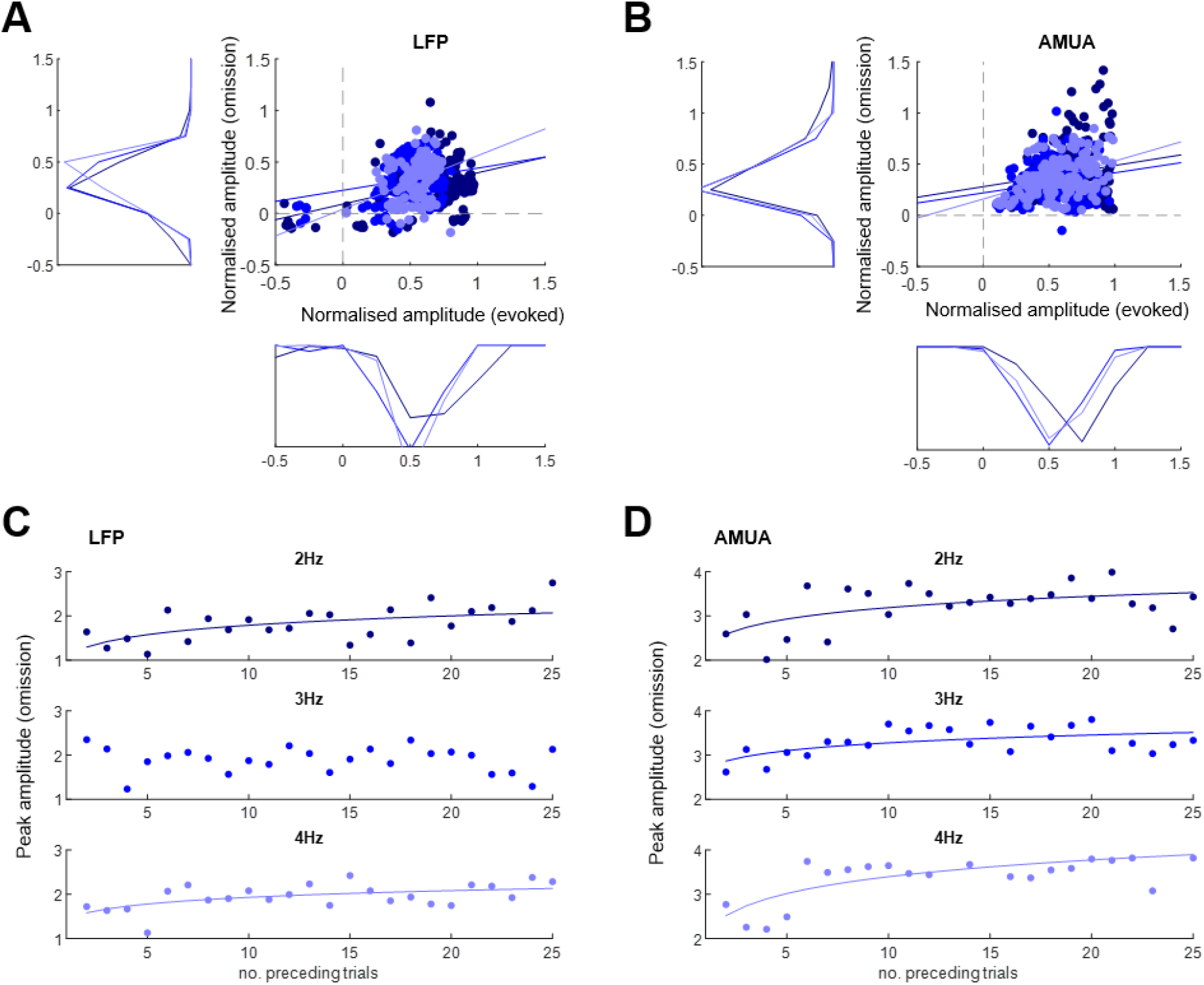
**(A)** Scatterplot and marginal histograms of peak LFP amplitude relationship between stimulus-evoked (X axis) and omission responses (Y axis) for each presentation rate (dark blue: 2 Hz, blue: 3 Hz, light blue: 4 Hz). Solid lines denote regression slopes. **(B)** Scatterplot and marginal histograms of peak AMUA amplitudes; legend as above. **(C)** Peak LFP amplitudes of omission responses as a function of the number of preceding noise bursts (dark blue: 2 Hz, blue: 3 Hz, light blue: 4 Hz). Solid lines denote significant regression slopes. **(D)** Peak AMUA amplitudes of omission responses as a function of the number of preceding noise bursts. Legend as above.

### 2.3 Buildup of omission responses over time

To characterise the modulation of omission responses as a function of the number of preceding noise bursts, we correlated peak amplitudes of omission responses with the number of preceding bursts of a particular rate, relative to the last omission. We log-transformed the number of preceding bursts to approximate exponential adaptation effects (Rajendran et al., 2017). This analysis revealed a significant increase of LFP omission responses as a function of the number of preceding bursts (Fig. 3C) for all but one burst rate (2 Hz: r = 0.5516, p = 0.0052; 3 Hz: r = −0.1448, p = 0.4996, n.s.; 4 Hz: r = 0.5292, p = 0.0078). All significant correlations survived Bonferroni correction for multiple comparisons. In the AMUA analysis (Fig. 3D), the correlation coefficients were significant for all analysed rates (2 Hz: r = 0.5306, p = 0.0076; 3 Hz: r = 0.5522, p = 0.0051; 4 Hz: r = 0.6998, p < 0.001). All significant correlations survived Bonferroni correction for multiple comparisons. Taken together, these findings are consistent with it taking time for the brain to build a model of the standard stimulus (noise burst), generate predictions, and signal errors (omission responses).

### 2.4 Laminar profile of omission-evoked responses

To analyse the laminar profile of omission-evoked responses, we transformed the LFP data into current source density (CSD) profiles to minimise the effect of volume conduction and obtain more precise estimates of local synaptic current flow (Lakatos et al., 2013). We tested whether the relative strength of omission and evoked responses shows any laminar differences, and whether these differences depend on the type of analysed responses (CSD vs. concomitant AMUA), by grouping responses into three layers by channel depth: superficial, intermediate, and deep (see Materials and Methods). To quantify the relative strength of omission responses, per channel, we calculated the omission response index (see Materials and Methods) by scaling the omission response amplitude to the average overall response amplitude (omission and evoked responses combined).

Overall, the omission response index was significantly higher for AMUA than for CSD (F_1,200_ = 8.53, p = 0.0046). Interestingly, we also found a significant interaction between signal type (AMUA vs. CSD) and laminar depth (F_2,200_ = 4.23, p = 0.0159). Given this interaction, we then compared the resulting omission response indices between response types (CSD vs. AMUA), separately for each laminar depth (Fig. 4). This analysis revealed that AMUA omission responses were relatively stronger than CSD omission responses in the superficial layer (two-sample t-test: t_71_ = 3.89, p < 0.001) and in the intermediate (input) layer (t_66_ = 3.21, p = 0.002), but not in the deep layer (t_82_ = 0.08, p = 0.938).

**Figure 4.**
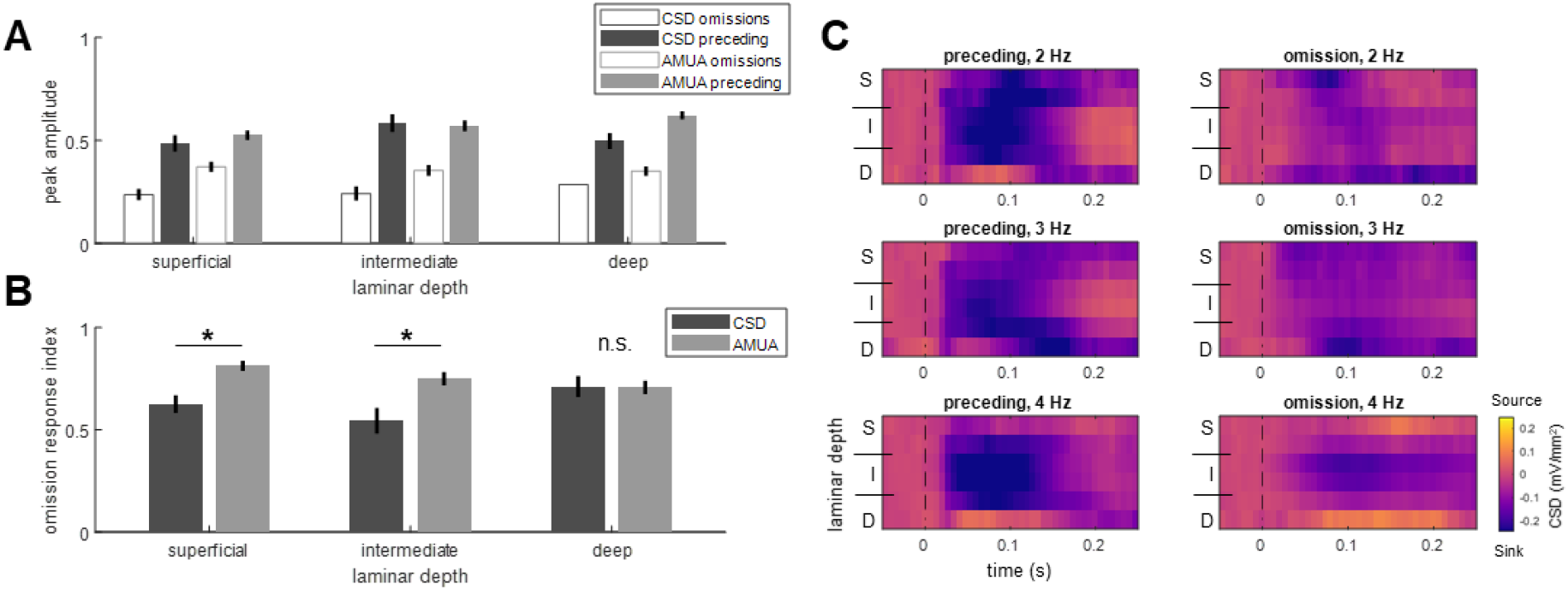
(A) Laminar specificity of omission responses: peak CSD and AMUA amplitudes plotted for stimulus-evoked (solid bars) and omission responses (empty bars) for CSD (cyan) and AMUA (magenta). Error bars represent SEM across channels. **(B)** Laminar specificity of the omission response index (see Materials and Methods) for CSD (cyan) and AMUA (magenta). Asterisk marks a significant difference between CSD and AMUA (p < 0.05, Bonferroni corrected across laminar depth). Error bars represent SEM across channels. **(C)** CSD maps per burst rate, plotted separately for stimulus-evoked and omission responses. Laminar depth: superficial (S), intermediate (I), deep (D) channels.

When analysing AMUA and CSD as a function of depth individually, we found no significant difference between layers in case of AMUA omission response indices (F_2,74_ = 1.82, p = 0.1693), but a significant difference between layers in case of CSD (F_2,135_ = 3.69, p = 0.0274). A post-hoc analysis of the latter finding showed a significant decrease of CSD omission response indices in the superficial vs. deep layers (t_94_ = 2.41, p = 0.0176), but not in the other pairwise comparisons (p > 0.13).

A visual inspection of CSD maps corresponding to omission responses (Fig. 4C) indicated that - at least for the fastest presentation rate - intermediate layers were characterised by activity sinks, while superficial layers were characterised by activity sources, suggesting that activity in these layers might predominantly correspond to output vs. input signals respectively.

### 2.4 Phase-locking of ongoing activity at omission time

Finally, to test whether – in addition to omission-evoked responses – omitted stimuli are associated with increased low-frequency synchronisation (entrainment) at the respective burst presentation rate, we analysed the phase locking value (PLV; see Materials and Methods), or intertrial phase coherence, of ongoing neural activity at the onset of an expected but omitted burst (Fig. 5). The PLV quantifies phase consistency across repeats - that is, does the waveform of the LFP at a particular frequency fluctuate in a similar way each time a burst is presented, or each time an omission is presented. We reasoned that neural entrainment to burst trains should result in increased PLV for the specific burst presentation rates, relative to other rates. Note that while PLV can be confounded by amplitude differences (with higher PLV obtained for signals with higher amplitudes), here we controlled for any differences in the power of neural activity by only selecting those channels which did not show differences in power between presentation rates (see Materials and Methods). On average, 16.33 channels per penetration were selected (SEM 3.55, corresponding to 25.51% ± 5.55% channels). This analysis revealed that PLV was modulated by the presentation rate (F_2,828_ = 2432.64, p < 0.001) and the frequency of neural activity (F_2,828_ = 4.85, p = 0.031), but not by the interaction between the two (F_4,828_ = 1.94, p = 0.141). In other words, we did not find frequency-specific PLV increases for the respective burst presentation rates. This suggests that any phase-locking (entrainment) of low-frequency activity is not specific to the presentation rate of preceding bursts, but rather scales non-specifically along the analysed frequency and presentation rate (higher PLV for lower presentation rates).

**Figure 5.**
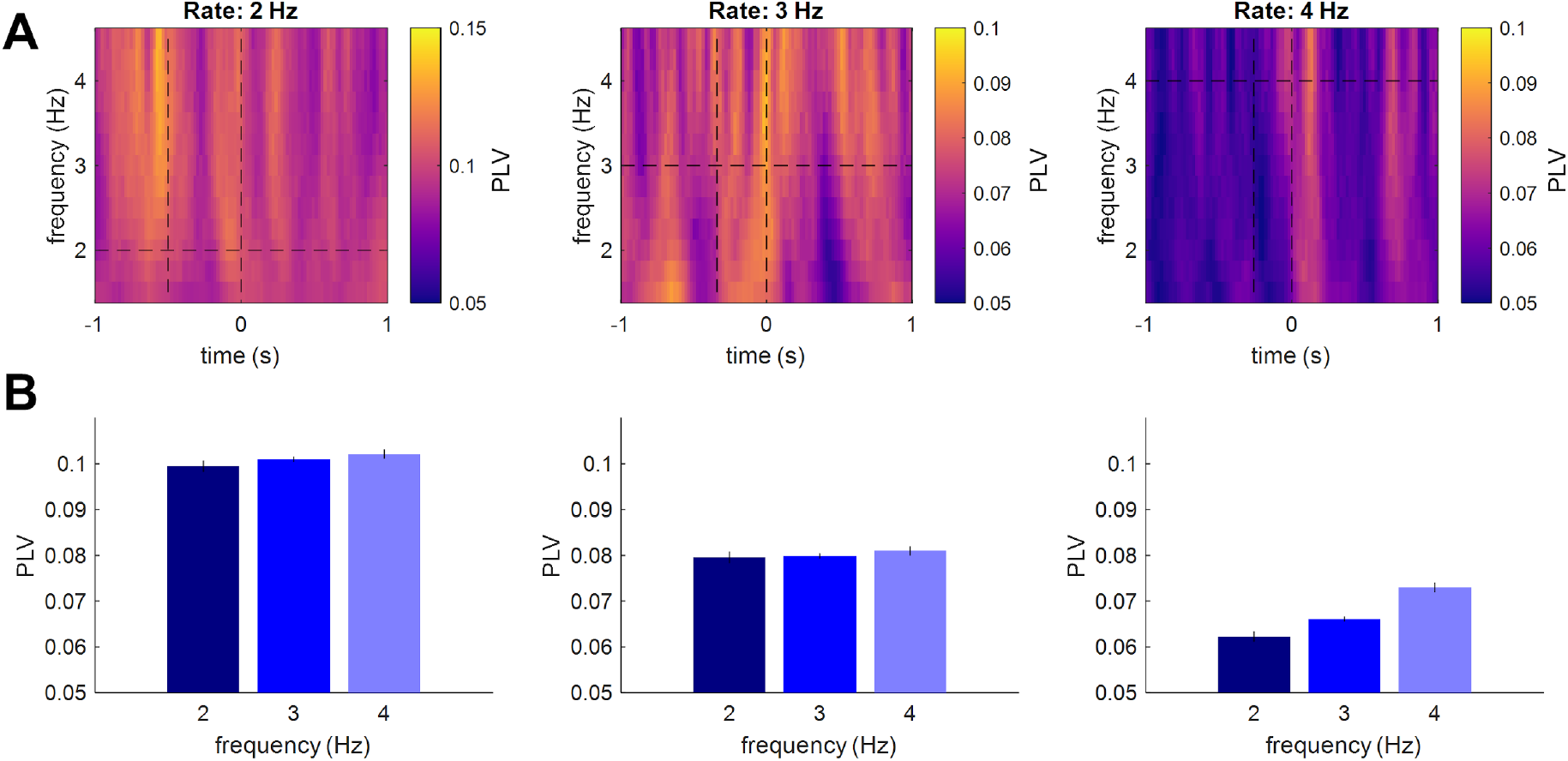
**(A)** Phase-locking value (PLV) time-frequency maps relative to the onset of an expected but omitted stimulus (right vertical dashed lines). Each panel represents blocks of rhythmic stimulation at a different presentation rate (left: 2 Hz, middle: 3 Hz, right: 4 Hz). Preceding stimulus latency is marked by the left vertical dashed line. Stimulus presentation frequency is marked by the horizontal dashed line. **(B)** PLV at expected but omitted stimulus onset. Different panels correspond to blocks of stimulation at different presentation rates. Different bars correspond to mean PLV values at omitted stimulus onset, estimated within a given block for different presentation rates (dark blue: 2 Hz, blue: 3 Hz, light blue: 4 Hz). Error bars represent SEM across channels.

In a similar analysis of PLV following sequence termination (one cycle after the last standard burst in the entire sequence), we observed a similar pattern of results. Both the main effects of the presentation rate (F_2,1044_ = 430.63, p < 0.001) and the frequency of neural activity (F_2,1044_ = 4.49, p = 0.036) were significant, but not the interaction between the two (F_4,1044_ = 1.76, p = 0.173). While these results should be interpreted with more caution given a lower number of trials per condition taken into the analysis (mean ± SEM: 27.94 ± 0.14) as compared to the remaining analyses, this suggests that no rate-specific entrainment was observed in our data either during stimulus omissions in the midst of a sequence, or following sequence termination.

## 3. DISCUSSION

We have identified robust omission responses in the auditory cortex of anaesthetised naive rats, suggesting that omission detection is an automatic brain function. These omission responses are temporally specific to the anticipated stimulus onset (but not to the onset of the preceding standard stimulus), consistent with their interpretation as representing a predictive signal rather than a non-specific offset response. However, they are observable only in the local field potentials and analogue multiunit activity, but not in spiking activity, suggesting that they might be mediated by mechanisms other than classical spiking activity of projection neurons. In the following, we discuss the response characteristics and functional significance of omission responses.

### 3.1 Omissions responses at the cellular level in the auditory cortex

The cellular-level omission responses that we observe in the auditory cortex bear similarities with more gross non-invasive recordings of omission responses from the auditory cortex of humans. The omission response we observed had lower amplitudes and longer latencies than stimulus-evoked responses, consistent with previous non-invasive results in humans (Andreou et al., 2015; Chennu et al., 2016; Todorovic et al., 2011). Non--invasive measurements of neural activity in humans have identified omission responses for a wide range of stimulus presentation rates, with ISIs as short as 200 ms (Yabe et al., 1997) or as long as 2 s (Busse and Woldorff, 2003), which spans the range we examined.

There have been a few studies examining omission response at a cellular level in the auditory cortex. Recent studies using calcium imaging in the auditory cortex of awake and anaesthetised naive mice have reported omission-related activity for long ISIs in the range of 2-4 s (Li et al., 2017; Wang et al., 2018), but they did not see them at 1 s ISI unlike in our results. The protocol for these calcium imaging studies was very different from ours, having no repeated omissions, the only omission being the end of a single stimulus train. Furthermore, neural activity measured using calcium imaging is characterised by much longer time constants relative to electrophysiological measurements, which makes it a suboptimal method of imaging time-courses of omission responses at faster presentation rates. Thus, our study is the first to show temporally specific omission responses in the auditory cortex of naive animals at presentation rates typical for most previous non-invasive human studies of neural activity (Chennu et al., 2016; Dercksen et al., 2020; Sanmiguel et al., 2013; Todorovic et al., 2011), and for repeated omissions. The omission signals observed in our study increased as a function of the (log) number of preceding noise bursts, suggesting that they might be modulated by the strength of predictions, based on the number of preceding standard stimuli (Rajendran et al., 2017). This gradual buildup of omission signals over time is consistent with the literature on mismatch responses following repetition suppression to consecutive standards (Auksztulewicz and Friston, 2016; Garrido et al., 2009a).

### 3.2 Omission responses under passive listening

While observing omission-related activity in naive animals suggests that it is a relatively automatic neural response, this conclusion is further reinforced by the fact that we measured neural activity from anaesthetised animals. This is constituent with previous reports that, in awake humans, omission responses can be observed under passive listening, with no attentional involvement (Chennu et al., 2016). To the best of our knowledge, no study so far has investigated omission responses in humans under anaesthesia or in disorders of consciousness. However, another type of neural response commonly associated with prediction error signals - namely, a mismatch response to deviant tones - is observed also under anaesthesia (Simpson et al., 2002) and in the vegetative state (Boly et al., 2011). In a previous study which compared omission-related activity in awake and anaesthetised mice (Li et al., 2017), omission responses were found to be only slightly more prevalent in the awake state (~21% neurons in the superficial layers of the auditory cortex showing omission responses in the awake state, compared to ~15% neurons under anaesthesia) - however, as noted above, this was at a slower rate than in our study. Taken together, our and previous results suggest that rudimentary temporal predictions, which result in omission responses to stimulus absence in a rhythmic context, are generated automatically, without the involvement of attention (behavioural relevance) or wakefulness.

### 3.3 Omission responses and entrainment

Previous studies have occasionally interpreted omission responses as resulting from neural entrainment (Lakatos et al., 2013; Li et al., 2017; Wang et al., 2018). According to the neural entrainment hypothesis, an influential model of the modulation of neural activity in sensory cortices by rhythmic stimulus presentation (Lakatos et al., 2019), isochronous sound delivery should gradually increase the phase locking of low-frequency activity in the auditory cortex, at a frequency specific to the stimulus presentation rate (Lakatos et al., 2013), resulting in strongest phase-locking at the exact times when omitted stimuli would be expected. In contrast, omission responses consist of broadband neural activity, including higher frequencies than the stimulus presentation rate, and peaking several hundreds of milliseconds after an expected stimulus is omitted. In this study we did not find evidence for entrainment, quantified as increased frequency-specific phase-locking at the onset of an expected or omitted stimulus. No low-frequency entrainment was observed when stimuli were omitted in the midst of a rhythmic sequence, as well as following sequence termination. While seemingly at odds with the previous study (Lakatos et al., 2013), it should be noted that the phase locking in the previous study was frequency-specific to the ISI only during sequence presentation; however, following sequence termination, entrainment was assessed as peaks in power rather than frequency-specific phase-locking. In another study of rhythmic temporal predictions (Breska and Deouell, 2017), PLV was averaged within a broader delta band (0.5 - 3 Hz) and shown to not differentiate between rhythmic (isochronous) and non-isochronous but temporally predictable sequences. Therefore, the hypothesis that neural entrainment should manifest as increased frequency-specific phase-locking also following stimulus omission has limited empirical evidence. Interestingly, a recent computational model based on an oscillator bank, while going beyond pure entrainment (sustained resonance to isochronous streams) and including recurrently connected neuronal populations to implement deviance detection, could only reproduce omission responses for ISIs below 200 ms (Chien et al., 2019). These results highlight the limitations of explaining temporal predictions in terms of oscillatory processes at slow time scales. In contrast, in our study, omission responses were observed in neural activity at a broad range of frequencies, including lower frequencies typically analysed as local field potentials (0.1 - 75 Hz) as well as high frequencies in the AMUA analysis (300 - 6000 Hz).

### 3.4 Omission responses along the auditory hierarchy

While omission responses have been reported at very early processing stages in the visual system (Schwartz et al., 2007), invasive and non-invasive studies suggest that auditory omissions are typically only found in the cortex (Andreou et al., 2015; Busse and Woldorff, 2003; Horváth et al., 2010; Karamürsel and Bullock, 2000; Li et al., 2017; Wang et al., 2018) exception of one study using intracellular recordings that found omission responses in the non-lemniscal thalamus (Gao et al., 2009). Omission responses have not been found in the inferior colliculus (Nishihara et al., 2014) or in the brainstem (Lehmann et al., 2016).

Based on our recordings in the auditory cortex, we observed that omission response amplitude was correlated with evoked response amplitude across channels, suggesting that the same areas that encode sounds might also signal sound omissions. Indeed, we found that channels were mostly sensitive to either sound alone (25-39% of analysed channels) or to both presented and omitted sounds (33-55% of analysed channels), and that very few channels (4-10%) were found to be sensitive to omissions alone. This is in contrast to a recent study that used relatively low-spatial resolution surface electrodes to measure broad local field potentials in human superior temporal gyrus, which contains higher auditory areas than we typically examined. This study reported neuronal populations that responded only to omitted sounds, but not to presented sounds (Fonken et al., 2019).

Taken together, these results are consistent with studies examining mismatch responses to deviant sounds, which found that mismatch-specific responses are more pronounced in hierarchically higher regions of the auditory pathway (Camalier et al., 2019; Casado-Román et al., 2020; Parras et al., 2017). Future studies should test whether a similar gradient might be observed for omission responses.

### 3.5 The implications of omission responses in field potentials but not spikes

One notable point for consideration is that while we see omission responses in the LFP and the AMUA, we do not see them in the single-unit or multi-unit activity. This has implications for the representation of prediction and prediction error. First we must consider what the LFP and AMUA represent. LFP is considered to represent the dendritic activity of neurons, likely from summed inputs but perhaps also from dendritic processing (Goense and Logothetis, 2008; Logothetis, 2002). The AMUA, while typically taken to represent the summed spiking activity, may in fact have substantial contributions from other neural potentials. For example, in auditory cortex the correlation is only ~0.6 between turning curves from spiking activity and the AMUA (Kayser et al., 2007), and also simulations suggest there is some power in massed excitatory postsynaptic potentials that is in the AMUA range, above 300Hz (Logothetis 2002). Indeed, recent work suggests that high spike rates in neocortex tend to correlate with field potential oscillations in the 50-180 Hz range, and less so with the lower range of LFPs or the higher range of the AMUA (Watson et al. 2017).

This suggests a number of non-exclusive possibilities. 1) Omission responses are present in the spike responses of neurons but only a small fraction of them. In our sample of 113 single- and multiunits, none showed a notable emission response, indicating that this fraction, if it exists, must be small. Furthermore, if this alone is the source of omission responses, it is hard to explain the size of the omission responses in the LFP and AMUA, being about half the size of the response to the presented stimuli. It is unclear how such a small fraction of neurons generate such a large LFP/AMUA response. One possibility is that this fraction of neurons is clustered at a particular layer in the auditory cortex that we did not sample sufficiently, however our sample of single and multi units spans the depth of the cortex and we did not see units responding to omissions at any depth. It is also possible that the fraction could grow under non-anaesthetised conditions. 2) Another possibility is that the neurons whose spikes signal the omission response are from regions outside the auditory cortex, and they synapse on auditory cortical neurons and alter their membrane potentials but this does not in turn impact the auditory cortical neurons’ spiking activity. This implies that the relevant resulting potentials are sub-threshold and perhaps somewhat isolated from the soma in a distant region of the neuron’s dendrites. Since cortical apical dendrites are electrotonically isolated from the site of spike generation at the axon hillock of the soma (Larkum et al., 2009, 1999), the omission responses could be in the potentials of these dendrites. 3) A third possibility is that the omission responses are calculated in the neuron by the summation of excitatory and inhibitory potentials, but that likewise this does not in turn impact the neurons’ spiking activity, again because it is sub-threshold and also perhaps isolated in the apical dendrites or other dendrites.

This paucity of neural spiking omission responses, accompanied by a strong field potential omission response, has implications for models of the cortex. It calls into question models which require many neurons whose spike-output signals prediction or prediction error, although it could be that such models could operate on timescales other than those that we examined. Instead, our findings may be somewhat congruent with recent modelling work which proposed that prediction error or related signals are represented in the apical dendrites of pyramidal neurons (Sacramento et al., 2017; Guerguiev et al., 2017; Richards & Lillicrap 2019). Given the complex hierarchical networks of the brain, how the brain assigns credit signals (such as prediction error) to the appropriate neurons and synapses to enable learning, without interfering with ongoing neural processing, is a key problem in neuroscience known as the credit assignment problem (Richards & Lillicrap 2019). It has been argued that the electrotonic isolation of the apical dendrites allows for the segregation of their proposed credit assignment calculations from the ongoing sensory integration at the soma and oblique and basal dendrites (Guerguiev et al., 2017; Richards & Lillicrap 2019).

## 4. MATERIALS AND METHODS

### 4.1 Auditory paradigm

Auditory stimuli were delivered binaurally using custom built in-ear headphones. The stimuli consisted of trains of broadband noise bursts presented at a fixed (isochronous) rate of 2, 3, or 4 Hz (Fig. 1B). Each noise burst was 25 ms long and 80 dB SPL. The noise bursts were embedded in a background of low-amplitude continuous white noise, at a burst-to-background ratio of 10 dB. Each stimulus train was 40 s long, with the first 36 s containing noise bursts and the last 4 s containing only low-amplitude background noise. In each train, a random subset of 5% bursts was omitted. In all but one experiment, each train started with at least 12 bursts with no omissions, and subsequent omissions were separated by at least 5 bursts. In the remaining experiment, omissions were implemented pseudo-randomly throughout the stimulus sequence (separated by at least 3 bursts). Per experiment, 90 trains were presented, divided into 9 blocks of 10 trains each. The noise burst rate did not change during each block. Between blocks, noise burst rate changed pseudorandomly (two consecutive blocks could not have the same burst rate). The placement of omissions in the sequence differed across stimulus trains and blocks. The burst rates were different across experiments.

### 4.2 Subjects and surgical procedures

All experimental procedures obtained approval and licences from the UK Home Office and followed legal requirements (ASPA 1986). The subjects were three female adult Lister hooded rats weighing 225 – 363 g (mean = 286 g) at the time of the experiment. Rats were anaesthetized with a mixture of 0.05 ml domitor (1mg/ml) and 0.1 ml ketamine (100mg/ml), administered intraperitoneally. To maintain anaesthesia, rats were infused continuously with a saline solution containing 16 μg/kg/h domitor, 4 mg/kg/h ketamine and 0.5 mg/kg/h torbugesic, at a rate of 1 ml/h. Body temperature was maintained with a heating pad at 36° ± 1° C. The depth of anaesthesia was controlled by regular testing of the absence of a toe pinch withdrawal reflex. The anaesthetised rats were placed in a stereotaxic frame with hollow ear bars set to fix the head for craniotomy. A craniotomy was performed with a centre at 4.7 mm caudal to bregma and 3.5 mm lateral to the midline. In two rats (rats 1 and 3), the craniotomy was performed over the right hemisphere; in one rat (rat 2), the craniotomy was performed over the left hemisphere.

### 4.3 Electrophysiological recordings and pre-processing

Electrophysiological data were recorded using a 64-channel silicon probe (Neuronexus Technologies, Ann Arbor, MI, USA) with 8 shanks each with 8 equally spaced electrodes along its length, forming a square grid pattern of 8×8 recording sites (electrode diameter: 175 μm^2^; distance between electrodes: 0.2 mm). Anatomical coordinates were used to position the probe over the auditory cortex. The probe was then inserted into the brain at a medio-lateral orientation until all recording sites were inside the brain. First, to verify that the recording sites were driven by sound stimulation, a search stimulus consisting of broadband noise bursts was played. Next, to check that channels contained signals from neuronal populations sensitive to acoustic frequency, frequency response areas (FRAs) were measured. Following these checks, experimental stimuli were presented binaurally via headphones at approximately 80 dB SPL. The stimulus sampling rate was set to 48828.125 Hz. Electrophysiological data were acquired at a sampling rate of 24414.0625 Hz using a TDT system 3 recording set-up (Tucker Davis Technologies). Across rats, data were recorded from 6 penetrations (rat 1: 1 penetration; rat 2: 3 penetrations; rat 3: 2 penetrations). In rats with multiple penetrations, the consecutive experiments were performed after moving the probe towards more rostral (rat 2) or dorsal (rat 3) sites by approx. 500 μm and repeating the search stimulus and FRA recordings.

Data traces (acquired in long segments of 40 s, corresponding to stimulus trains) were filtered off-line using 7th-order two-pass Butterworth filters: a notch filter (49-51 Hz) to remove line noise, and a high-pass filter (cut-off frequency: 0.1 Hz) to remove low-frequency drifts. Data were then epoched into shorter segments, corresponding to stimulus omissions (from −100 ms to 250 ms relative to the onset of expected but omitted stimuli) and immediately preceding noise bursts (from −100 ms to 250 ms relative to burst onset). The resulting number of omission and preceding burst epochs per penetration (mean ± SD) was 89 ± 4.89 for the 2 Hz burst rate; 165.66 ± 18.78 for the 3 Hz burst rate; and 232.16 ± 0.41 for the 4 Hz burst rate.

### 4.4 Data analysis

#### 4.4.1 Amplitudes and latencies of omission-evoked responses

Short segments were analysed in three ways, to obtain measures of local field potentials (LFP), analog multiunit activity (AMUA), and single- and multiunit spiking activity. In the LFP analysis, low-frequency signals were derived from original data by low-pass filtering each short segment using a 3rd order two-pass Butterworth filter (cut-off frequency: 75 Hz) and downsampling to 150 Hz. In the AMUA analysis, data were band-pass filtered using a 3rd order two-pass Butterworth filter between 300 and 6000 Hz (Lakatos et al., 2020). Then the analytic envelope (calculated using a 2-tap FIR filter) of the band-pass data was downsampled to 150 Hz. In both analyses, the epoched traces were normalised by z-scoring each trace relative to the pre-stimulus baseline (during the 70 dB background noise). To remove outliers, we calculated a standard deviation of the voltage fluctuation in each trial (SDi) and rejected trials with SDi beyond the median ± 3 SD of all SDi values.

To test whether single channels show noise-burst-evoked activity, single-trial amplitudes were averaged over time (from 0 ms to 250 ms relative to burst onset), pooled over burst rates, and (since the data were already normalised to the pre-stimulus baseline) entered into a one-sample t-test for each channel. Similarly, to test whether single channels show omission-evoked activity, amplitudes were averaged over time (from 0 ms to 250 ms relative to expected but omitted burst onset), pooled over burst rates, and subjected to one-sample t-tests. The resulting p-values were corrected for multiple comparisons using a false-discovery rate p_FDR_ < 0.05 (Benjamini and Hochberg, 1995). Only those channels showing both significant burst- and omission-evoked responses were entered into subsequent analyses (however, in a control analysis, we also analysed data from all channels, without any selection criteria). Single-trial data from the selected channels were averaged per penetration, channel, burst rate (2, 3, and 4 Hz), and stimulus type (burst vs. omission). Average traces were used to extract peak amplitudes and latencies.

Omission-evoked responses were compared with burst-evoked responses using mixed-effects modelling. In separate analyses, we compared (1) peak amplitudes, (2) peak latencies, and (3) entire time courses of burst-evoked and omission-evoked responses. In all cases, single-channel data were entered into a mixed-effects model with two fixed-effects factors (stimulus type: burst vs. omission; burst rate: 2, 3, and 4 Hz) and one random-effects factor (penetration). In analysing the time-courses, statistical tests were performed per time point and corrected for multiple comparisons using a false-discovery rate p_FDR_ < 0.05. In a control analysis, to test for the possibility that omission-related activity is due to random noise fluctuations in the post-omission time window rather than to true omission responses, we repeated the analysis described above but after shuffling single-trial data over time. We reasoned that if omission responses are due to noise fluctuations, shuffling data over time should not affect the peak amplitudes. Conversely, if omission responses reflect neural activity following an expected but omitted stimulus, shuffling data will abolish any omission-locked activity peaks.

To test whether omission responses build up over time (as a function of the number of preceding noise bursts), for each trial and analysed channel we extracted the peak amplitude of the omission response, averaged these peak amplitudes across channels, and correlated them with the log number of preceding noise bursts to model exponential decay (Rajendran et al., 2017). The Pearson correlation analysis was conducted separately for each signal type (LFP, AMUA) and burst rate (2, 3, and 4 Hz). Significance of correlation coefficients was Bonferroni-corrected for multiple comparisons.

Finally, in the analysis of spiking activity, we performed offline spike sorting using the expectation-maximization algorithm Klustakwik (Kadir et al., 2014) followed by manual post-processing using the Klustaviewa toolbox (Cortical Processing Lab, University College London). The algorithm returns two types of clusters of spikes - one putatively originating from a single neuron (termed a single unit, n = 79 across 6 penetrations) and one putatively originating from a small population of neurons (termed a multiunit, n = 280) near a recording site. Firing rate time series were calculated by binning spike times into 10 ms bins, resulting in peri-stimulus time histograms (PSTHs) sampled at 100 Hz. Only those single units and multiunits that were reliably driven by stimuli (noise bursts) were included in further analysis. In order to quantify firing reliability, we used a noise power to signal power metric (Sahani & Linden, 2003), which characterises the repeatability of neural response patterns for multiple presentations of the same stimulus. Neural responses to the first 250 ms of all 3 burst rates (2, 3, and 4 Hz) were concatenated for this analysis. Following previous studies (Rabinowitz et al., 2012; Rajendran et al., 2020), only those units showing a noise power ratio higher than 40 were included in the analysis, amounting to 43 single units (54.43%) and 70 multiunits (25%).

PSTHs were analysed to test for significant omission-evoked responses. Baseline spontaneous firing rate (SFR) was quantified as the average firing rate during the 50 ms preceding noise burst onset. To generate a null distribution, 1000 simulated PSTHs were calculated using a Poisson model assuming a constant firing rate equal to the SFR (Parras et al., 2017). For both actual and simulated PSTHs, response amplitudes were baseline-corrected by subtracting the SFR. Post-omission baseline-corrected PSTHs were tested for statistical significance by calculating the p-value of the actual PSTH as p = (k + 1) / (N + 1), where k is the count of simulated PSTHs for which the root mean square (RMS) over the post-stimulus period (0-250 ms, averaged across burst rates) was greater than or equal to the RMS of the actual PSTH, and N = 1000 simulations. This procedure could yield a minimum p ≈ 0.001. The resulting p-values were corrected for multiple comparisons using a false-discovery rate p_FDR_ < 0.05 (Benjamini and Hochberg, 1995).

#### 4.4.2 Laminar profile of omission-evoked responses

To test whether omission responses preferentially engage superficial, intermediate, or deep cortical layers, and whether the laminar profile depends on the type of responses (higher-frequency / AMUA vs. lower-frequency / LFP), we analysed peak amplitudes at electrodes which yielded significant omission responses of both types (AMUA and LFP). In order to increase the laminar resolution of the LFP signals, they were converted to current source density (CSD) estimates by calculating the second spatial derivative over channels. Given that each shank contained 8 electrodes, this procedure resulted in 6 CSD estimates as a function of channel depth. To make AMUA and CSD data more comparable, we therefore removed the edge channels from AMUA analysis. Data were pooled across penetrations, and electrode shanks into three groups of channels: superficial (channels 2-3 of each shank), intermediate (channels 4-5), and deep (channels 6-7). The same criteria were applied to assigning single units and multiunits (SUA, MUA) to different layers, except for edge channels being included (superficial: channels 1-3; intermediate: channels 4-5; deep: channels 6-8). To summarise the relative strength of omission responses, we calculated the omission response index of each channel (i.e., the peak amplitude of the omission response divided by the average peak amplitude of the omission and burst-evoked response). The resulting omission response index was always positive, lower than 1 if the omission response was weaker than the burst-evoked response, and higher than 1 if the omission response was stronger than the burst-evoked response. Omission response indices were compared between response types (AMUA vs. CSD) and channel groups (superficial, intermediate, deep) in a 2×3 ANOVA across channels. Post-hoc two-sample t-tests (separate for each channel group) were corrected for multiple comparisons using Bonferroni correction.

#### 4.4.3 Phase-locking of ongoing activity at omission time

Finally, to test whether, in addition to omission-evoked responses, in the human EEG data omissions are also associated with increased entrainment (i.e., the intertrial phase consistency of ongoing oscillations at a low frequency of interest) to the presentation rate of preceding noise bursts, we analysed the phase-locking value (PLV) (Lachaux et al., 1999) of ongoing activity at the time of the expected but omitted burst. Hypothetically, one might see no increase in response around the omitted stimulus, but the phase of the ongoing activity could still lock to the timing of the omission. To this end, we computed instantaneous power and phase of ongoing activity in the 1.5-4.5 Hz range (in 0.25 Hz steps) at each time point from −1000 to 1000 ms (in 20 ms steps) relative to the expected but omitted burst onset using a one-cycle Morlet wavelet transform. A single cycle was chosen to minimise the influence of activity evoked by preceding and subsequent bursts on the phase estimates at the expected onset of an omitted burst. The extracted phase values were used to calculate PLV for each time-frequency point according to the equation below, where φ(n) is the instantaneous phase estimate calculated for each n of N trials and 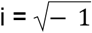:

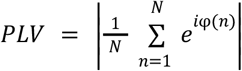

To test whether the presentation rate of preceding noise bursts modulates PLV at the expected onset of an omitted burst, we averaged PLV estimates within a half-cycle of a given frequency (e.g., for 2 Hz, within −250 to 250 ms relative to the omitted burst). Since PLV effects can be biased by differences in power between conditions (van Diepen and Mazaheri, 2018), we first analysed electrodes with respect to instantaneous power, selecting only those electrodes with no significant differences in power between presentation rates. To this end, per electrode, single-trial power estimates at the expected but omitted burst onset were extracted for 3 wavelet frequencies (2, 3, and 4 Hz), averaged within a half-cycle of a given frequency, and entered into a two-factor ANOVA (factors: wavelet frequency and presentation rate; both at 3 levels: 2, 3, and 4 Hz). Only those electrodes for which power estimates did not yield either a significant interaction effect of presentation rate and wavelet frequency, or a significant main effect of presentation rate (both p > 0.05), were used in subsequent PLV analysis. Here, the PLV estimates at the expected but omitted burst onset were entered into a three-factor mixed-effects analysis (fixed factors: wavelet frequency and presentation rate; random factor: penetration).

Finally, aiming at replicating the previous findings of temporally specific echo responses following sequence termination, we calculated PLV one cycle after the end of the sequence, using identical time-frequency decomposition methods, channel selection, and statistical tests as above.

## SUPPLEMENTARY FIGURES

**Figure S1.**
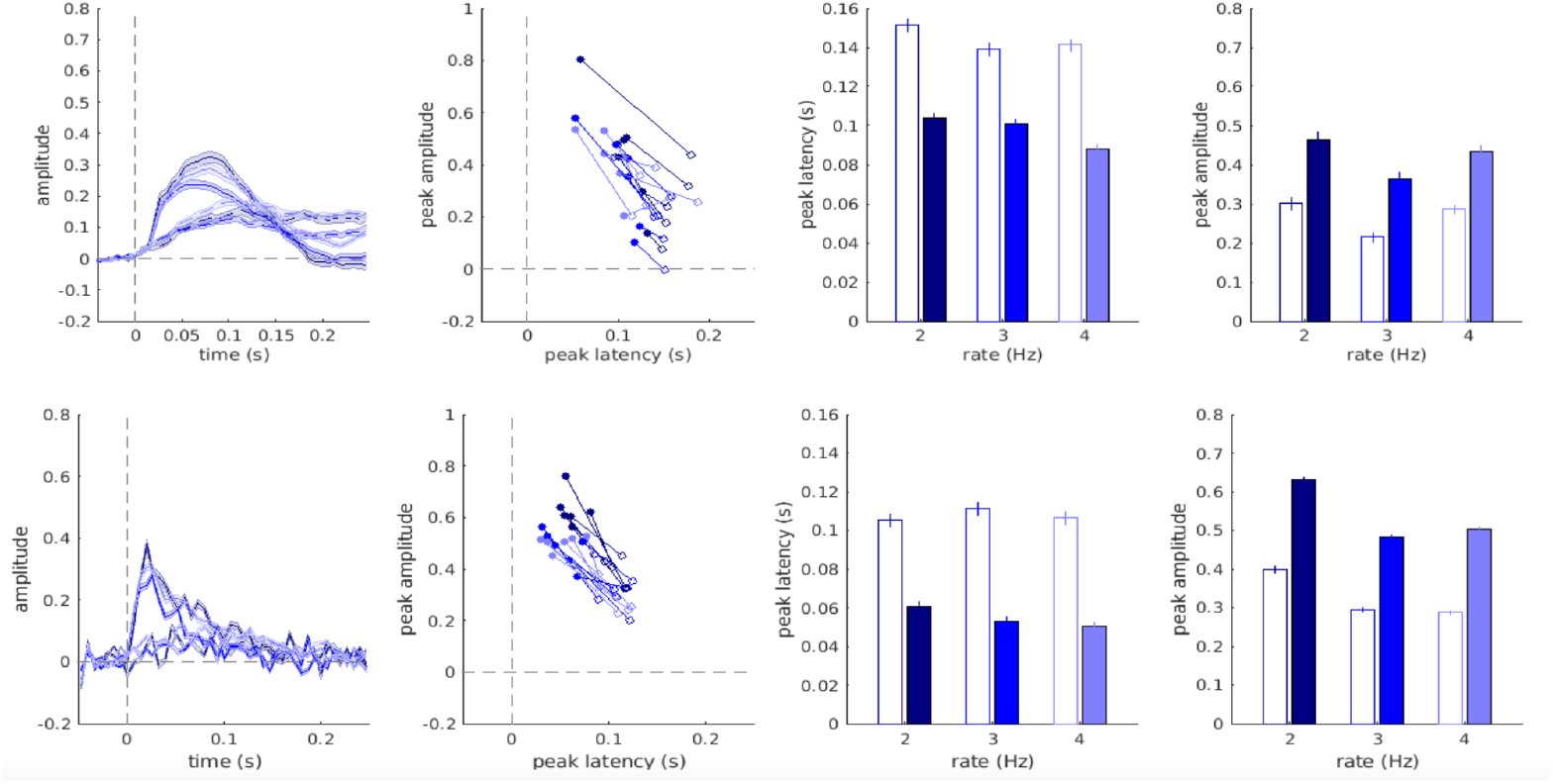
LFP (upper panels) and AMUA (lower panels) activity analysed for all channels, with no channel selection criteria. Figure legend as in Fig. 2A-H.

**Figure S2.**
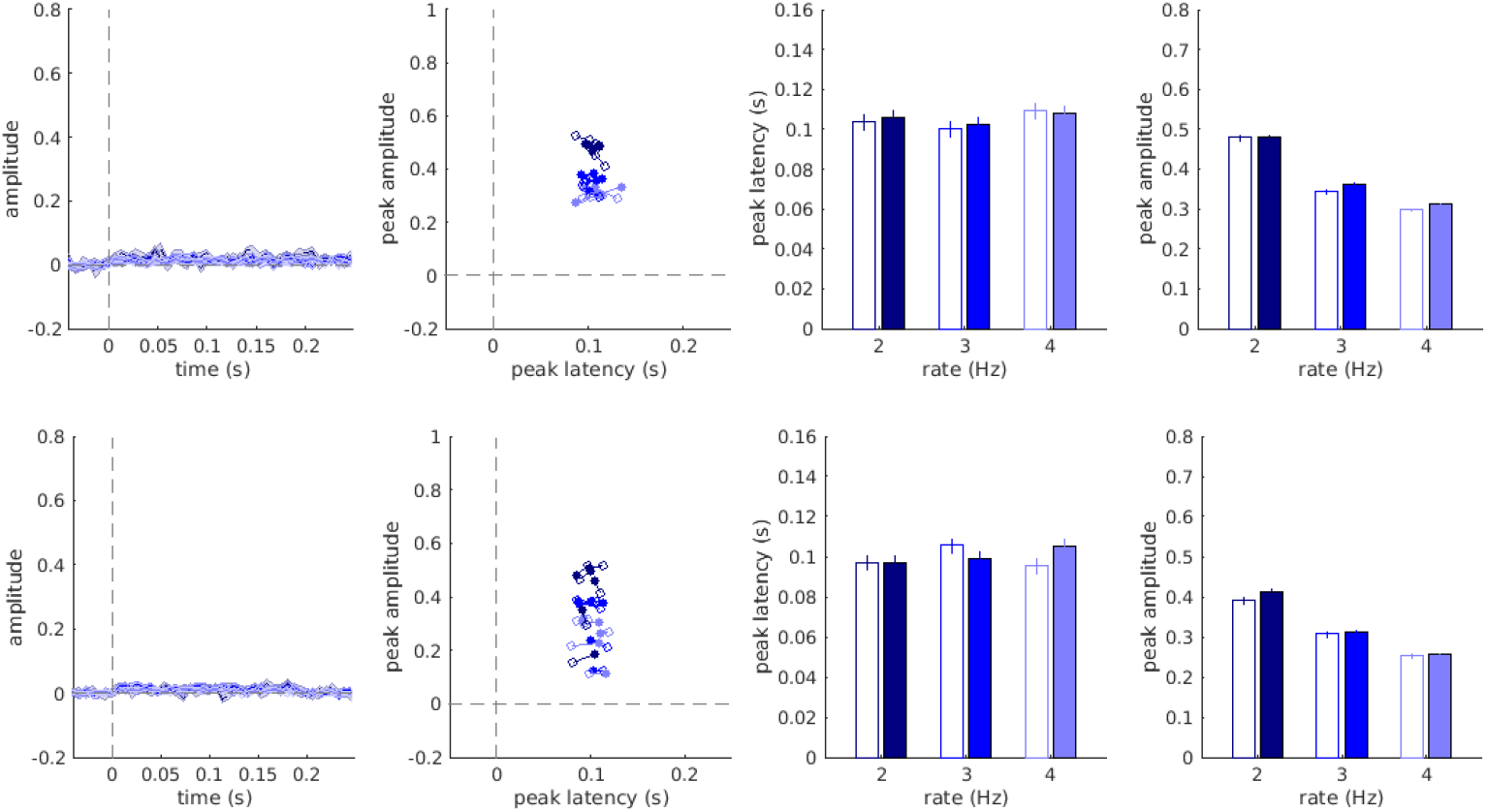
LFP (upper panels) and AMUA (lower panels) activity analysed for time-shuffled data. Figure legend as in Fig. 2A-H.

## ACKNOWLEDGEMENTS

This work has been supported by the European Commission’s Marie Skłodowska-Curie Global Fellowship (750459 to R.A.), the Hong Kong General Research Fund (11100518 to R.A., N.H. and J.S.), a grant from European Community/Hong Kong Research Grants Council Joint Research Scheme (9051402 to R.A. and J.S.), and the Wellcome Trust (WT09975MA, to V.R.). NH was supported by the Wellcome Trust (grant no. WT108369/2015/Z). We would like to thank Kerry Walker for helpful discussions.

